# Microbiome–host systems interactions: Protective effects of propionate upon the blood–brain barrier

**DOI:** 10.1101/170548

**Authors:** Lesley Hoyles, Tom Snelling, Umm-Kulthum Umlai, Jeremy K. Nicholson, Simon R. Carding, Robert C. Glen, Simon McArthur

## Abstract

**Background:** Gut microbiota composition and function are symbiotically linked with host health, and altered in metabolic, inflammatory and neurodegenerative disorders. Three recognized mechanisms exist by which the microbiome influences the gut-brain axis: modification of autonomic/sensorimotor connections, immune activation, and neuroendocrine pathway regulation. We hypothesized interactions between circulating gut–derived microbial metabolites and the blood–brain barrier (BBB) also contribute to the gut–brain axis. Propionate, produced from dietary substrates by colonic bacteria, stimulates intestinal gluconeogenesis and is associated with reduced stress behaviours, but its potential endocrine role has not been addressed.

**Results:** After demonstrating expression of the propionate receptor FFAR3 on human brain endothelium, we examined the impact of a physiologically relevant propionate concentration (1 μM) on BBB properties *in vitro*. Propionate inhibited pathways associated with non-specific microbial infections via a CD14-dependent mechanism, suppressed expression of LRP-1 and protected the BBB from oxidative stress via NRF2 (NFE2L2) signaling.

**Conclusions:** Together, these results suggest gut-derived microbial metabolites interact with the BBB, representing a fourth facet of the gut–brain axis that warrants further attention.

## Background

The human body plays host to, and exists in symbiosis with, a significant number of microbial communities, including those of the skin, oral and vaginal mucosae and, most prominently, the gut [1]. This relationship extends beyond simple commensalism to represent a major regulatory influence in health and disease, with changes in abundance of members of the faecal microbiota having been associated with numerous pathologies, including diabetes, hepatic diseases, inflammatory bowel disease, viral infections and neurodegenerative disorders [2–8]. Metagenomic studies have revealed reductions in microbial gene richness and changes in functional capabilities of the faecal microbiota to be signatures of obesity, liver disease and type II diabetes, and that these can be modified by dietary interventions [9,10]. The gut microbiome harbours 150 times more genes than the human genome, significantly increasing the repertoire of functional genes available to the host and contributing to the harvesting of energy from food [11].

The primary form of communication within the gut microbe–human super-system is metabolic, but our understanding of the details of the cross-signalling pathways involved is limited. It is clear, however, that gut-derived microbial metabolites and products such as lipopolysaccharide (LPS) can influence human health both in the intestine and systemically [12,13], with reported effects ranging from mediation of xenobiotic toxicity [14], through modification of the risk of preterm birth [15] to induction of epigenetic programming in multiple host tissues [16,17]. A major aspect of microbe–host systems-level communication that is receiving increased attention is the influence the gut microbiota exerts upon the central nervous system (CNS), the so-called ‘gut–brain axis’ [18].

The existence of gut–brain communication is supported by a number of animal and human studies, although the underlying mechanisms are not always well defined. Behavioural analysis of antibiotic-treated or germ-free rodents reveals alterations in both stress responsiveness [19] and anxiety [20–22], although in germ-free models these findings are complicated by the life-long absence of gut microbes and possible consequent developmental alterations. Nonetheless, gut-microbe-depleted animals have been shown to exhibit changes in serotonergic and glutamatergic neuronal signalling [20] and expression of brain-derived neurotrophic factor (BDNF) within the limbic system [22,23], providing a molecular correlate for behavioural changes.

Links between the gut microbiota and brain function have been identified in studies of humans with autism spectrum disorders (ASD) and attention-deficit hyperactivity disorder (ADHD). Altered microbial profiles have been identified in children with ASD [24–26], and oral treatment of autistic children with the non-absorbed, broad-spectrum antibiotic vancomycin – effectively suppressing the gut microbiota – led to a regression in autistic behavioural characteristics that was reversed upon antibiotic discontinuation [27]. Similarly, a small-scale intervention study has suggested not only a link between lower counts of faecal *Bifidobacterium* species at six months and increased incidence of ADHD at 13 years, but also that early probiotic treatment lessens the risk of ADHD development [28].

A number of unresolved questions remain as to the mechanism(s) of communication between the gut microbiota and the brain, but three major pathways have been proposed: direct modification of vagal or sympathetic sensorimotor function [29], inflammatory/immune activity [30] and neuroendocrine crosstalk [31]. While research in this field has focussed most heavily on direct neural modulation and inflammatory signalling, the potential role of circulating gut microbe-derived metabolites has been relatively underexplored. Communication with and across the blood–brain barrier (BBB), the primary interface between the circulation and the CNS, may therefore represent a significant mechanism allowing the gut microbiota to influence brain function.

There is accumulating evidence that the gut microbiota can affect the integrity of the BBB, with both broad-spectrum-antibiotic-treated and germ-free mice exhibiting considerably enhanced barrier permeability and dysregulation of inter-endothelial cell tight junctions [32,33]. Importantly, these impairments can be reversed upon conventionalisation. The mechanism(s) by which gut microbes exert their influence are unclear, but changes to brain chemistry induced by alteration of the gut microbiota can occur independently of vagal or sympathetic neural pathways and in the absence of any immune response, strongly suggesting at least a contributory role for soluble gut-derived microbial metabolites [22].

In particular, data highlight a potential role for short-chain fatty acids (SCFAs) as key microbial mediators in the gut–brain axis. SCFAs are principally produced by the fermentation of complex plant-based polysaccharides by gut bacteria and are potent bioactive molecules; stimulating colonic blood flow and upper-gut motility, influencing H2O and NaCl uptake, providing energy for colonocytes, enhancing satiety and positively influencing metabolic health in obese and diabetic individuals [34–36]. Of the SCFAs, acetate is produced in the greatest quantity as a result of fermentation in the large intestine, followed by propionate and butyrate [37]. Over 95 % of SCFAs produced are absorbed within the colon with virtually none appearing in the urine or faeces [35,38]. However, all three metabolites are detectable in the peripheral blood of healthy individuals (http://www.hmdb.ca: acetate, 22–42 μM; propionate, 0.9–1.2 μM; butyrate, 0.3–1.5 μM). SCFAs activate members of the free fatty acid receptor (FFAR) family of G protein coupled receptors; acetate, propionate and butyrate have affinity in the low millimolar to high micromolar range for FFAR2; propionate and butyrate have mid to low micromolar affinity for FFAR3 [39].

The majority of studies looking at the role of SCFAs in the gut–brain axis have focused on butyrate [40], with relatively few investigating propionate despite its similar plasma concentration and receptor affinity. Propionate is a highly potent FFAR3 agonist for its size (agonist activity GTPγS pEC_50_ (E_max_) 3.9-5.7(100%)) and has close to optimal ligand efficiency (-Δ*G*=1.26 kcal mol^-1^ atom^-1^) for this receptor [41]. While propionate has been shown to stimulate intestinal gluconeogenesis through direct stimulation of enteric–CNS pathways [42], and increased intestinal propionate has been associated with reduced stress behaviours [43] and reward pathway activity [44] in mice and humans, respectively, its potential role as an endocrine mediator in the gut–brain axis has not been addressed. Given the presence of FFAR3 on endothelial cells [45], we hypothesised that propionate targeting of the endothelium of the BBB would represent an additional facet of the gut–brain axis. We used a systems approach to test this proposal, performing an unbiased study of the transcriptomic effects of exposure to physiological levels of propionate upon the BBB, modelled by the immortalised human cerebromicrovascular endothelial cell line hCMEC/D3, accompanied by *in vitro* validation of identified pathway responses.

## Results

### Microarray analyses

Following initial confirmation of the expression of FFAR3 in human brain endothelium (**Fig. 1a**) and on hCMEC/D3 cells (**Fig. 1b**), we investigated the effect of exposure of hCMEC/D3 monolayers to 1 μM propionate for 24 h. Such treatment had a significant (*P*_FDR_ < 0.1) effect on the expression of 1136 genes: 553 upregulated, 583 downregulated (**Fig. 1c**). Initially, we used SPIA with all the significantly differentially expressed genes to identify KEGG signalling pathways inhibited and activated in the presence of propionate. Protein processing in the endoplasmic reticulum and RNA transport were activated upon exposure of cells to propionate, which was unsurprising given gene expression had been induced. A number of pathways associated with nonspecific microbial infections (Gram-negative bacteria, viral) were inhibited by propionate (**Fig. 1d**), as were the cytosolic DNA-sensing pathway (upregulated by pathogen DNA during microbial infections, triggering innate immune signalling [46]), the NFκB signalling pathway and the Toll-like receptor signalling pathway. Of the 19309 genes we examined on the array, 203 of the 224 genes known to be associated with the BBB were detected (**Supplementary Table 1**). Eleven of these were significantly differentially expressed, with the majority being associated with the inflammatory response.

**Fig. 1:**
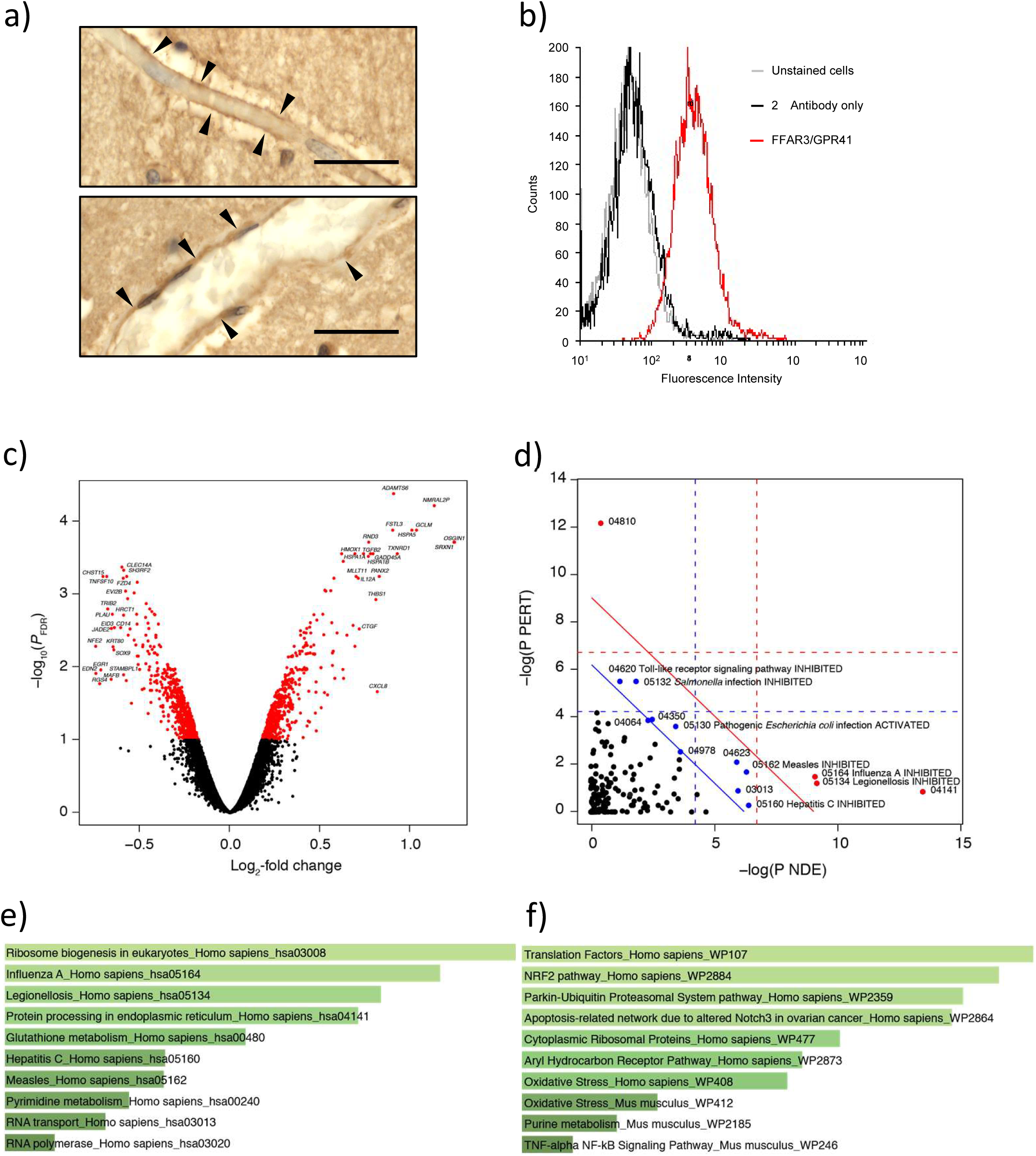
Effects on gene expression of exposure of the hCMEC/D3 cell line to propionate (1 μM, 24 h). (a) Representative images of FFAR3 immunoreactivity within endothelial cells of capillaries (i) and larger post-capillary (ii) blood vessels in control human brains *post mortem*; scale bar 20 μm, sections are 5 μm thick; images are representative of five independent cases, areas of particular immunoreactivity are highlighted by black arrowheads. (b) Surface expression of FFAR3/GPR41 by hCMEC/D3 cells (grey line, unstained cells, black line secondary antibody control, red line FFAR3), data are representative of three independent experiments. (c) Volcano plot showing significantly (*P*_FDR_ < 0.1, red dots) differentially expressed genes. The top 20 up- and down-regulated genes are labelled. (d) SPIA evidence plot for the 1136 significantly differentially expressed genes. Only those human KEGG pathways associated with non-specific microbial infections are labelled. The pathways at the right of the red oblique line are significant (*P* < 0.2) after Bonferroni correction of the global *P* values, pG, obtained by combining the pPERT and pNDE using the normal inversion method. The pathways at the right of the blue oblique line are significant (*P* < 0.2) after a FDR correction of the global *P* values, pG. 04810, Regulation of actin cytoskeleton (inhibited); 04064, NF-kappa B signaling pathway (inhibited); 04978, Mineral absorption (inhibited); 03013, RNA transport (activated); 04141, Protein processing in endoplasmic reticulum (activated); 04350, TGF-beta signaling pathway (activated); 04623, Cytosolic DNA-sensing pathway (inhibited). (e) Association of all significantly differentially expressed genes (*n* = 1136) with KEGG pathways, Enrichr. (f) Association of all significantly upregulated genes (*n* = 553) with WikiPathways, Enrichr. (e, f) The lighter in colour and the longer the bars, the more significant the result is. Significance of data was determined using rank-based ranking; only the top 10 results are shown in each case.

Enrichr [47,48] was used to examine KEGG pathways significantly associated with the list of significantly differentially expressed genes. All 1136 significantly differentially expressed genes mapped to Enrichr. As with SPIA, the genes were associated with KEGG pathways implicated in non-specific microbial infections, and RNA- and endoplasmic reticulum-associated processes (**Fig. 1e**).

WikiPathways analysis (Enrichr) of all the significantly differentially expressed genes highlighted responses to oxidative stress being associated with propionate treatment (not shown). Closer examination of the data demonstrated this was linked to NRF2 (NFE2L2) signaling, with the significantly upregulated genes closely associated with oxidative stress responses (**Fig. 1f**).

### Pathway validation

Transcriptomic analysis identified two particular clusters of pathways as being regulated by propionate treatment: those involved in the non-specific inflammatory response to microbial products (**Fig. 1d, e**) and those involved in the response to oxidative stress (**Fig. 1f**). We, therefore, sought to validate these responses in an *in vitro* model of the BBB.

### TLR-specific pathway

Inhibition of the TLR-specific pathway by propionate suggests this metabolite may have a protective role against exposure of the BBB to bacterial lipopolysaccharide (LPS), derived from the cell walls of Gram-negative bacteria. In accord with this hypothesis, exposure of hCMEC/D3 monolayers for 12 h to propionate at physiological concentrations (1 μM) was able to significantly attenuate the permeabilising effects of exposure to *Escherichia coli* O111:B4 LPS (subsequent 12 h stimulation, 50 ng/ml), measured both through paracellular permeability to a 70 kDa FITC-conjugated dextran tracer (**Fig. 2a**) and trans-endothelial electrical resistance (**Fig. 2b**). To determine the specificity of these effects for propionate, we investigated the actions of the closely related SCFAs acetate and butyrate. While physiologically relevant circulating concentrations of butyrate (1 μM) replicated the effects of propionate on both trans-endothelial electrical resistance and paracellular tracer permeability, this was not the case for acetate (65 μM) (**Fig. 2a-b**).

**Fig. 2:**
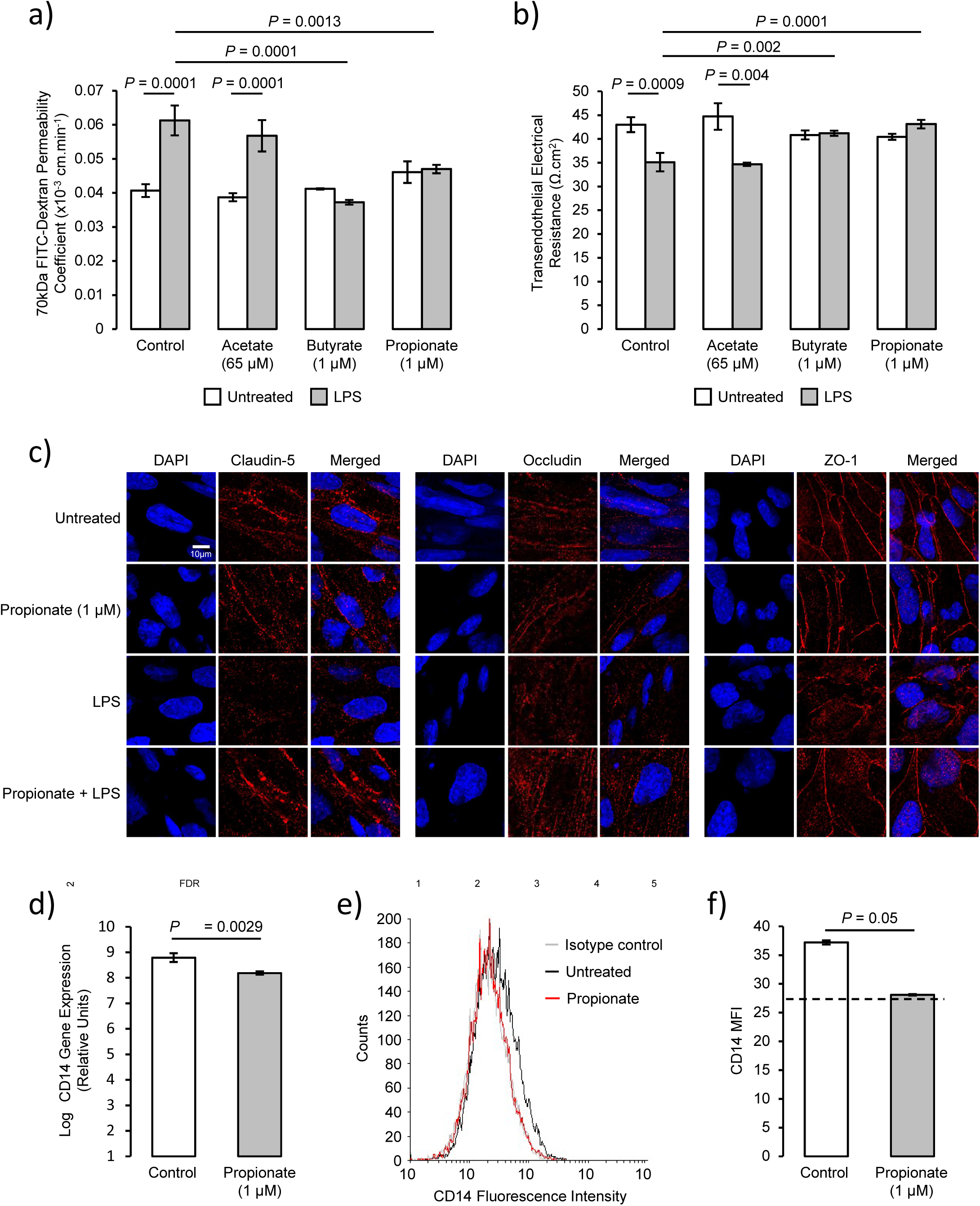
Protective effects of propionate against LPS-induced barrier disruption. (a) Assessment of the paracellular permeability of hCMEC/D3 monolayers to 70 kDa FITC-dextran following treatment for 24 h with 65 μM acetate, 1 μM butyrate or 1 μM propionate, with or without inclusion of 50 ng/ml LPS for the last 12 h of incubation; data are mean ± SEM, *n* = 3 independent experiments. (b) Trans-endothelial electrical resistance of hCMEC/D3 monolayers following treatment for 24 h with 65 μM acetate, 1 μM butyrate or 1 μM propionate, with or without inclusion of 50 ng/ml LPS for the last 12 h of incubation; data are mean ± SEM, *n* = 3 independent experiments. (c) Confocal microscopic analysis of expression of the tight junction components claudin-5, occludin and zona occludens-1 (ZO-1) in hCMEC/D3 cells following treatment for 24 h with 1 μM propionate, with or without inclusion of 50 ng/ml LPS for the last 12 h of incubation. Scale bar (10 μm) applies to all images. Images are representative of at least three independent experiments. (d) Expression of *CD14* mRNA in control and propionate-treated (1 μM; 24 h) hCMEC/D3 cells according to microarray data (data are mean ± SEM, *n* = 3). (e) Surface expression of CD14 protein on control and propionate-treated hCMEC/D3 cells (grey line, unstained cells, black line secondary antibody control, red line FFAR3), data are representative of three independent experiments. (f) Median fluorescence intensity of surface expression of CD14 protein on control and propionate-treated hCMEC/D3 cells, dashed line indicates isotype control fluorescence intensity; data are mean ± SEM, *n*=3 independent experiments.

Circulating concentrations of propionate are approximately 1 μM at rest, but these may be expected to increase following consumption of, for example, a meal containing high levels of fermentable fibre [1], consequently we examined the effects of 10 μM and 100 μM propionate upon the response of hCMEC/D3 monolayers to LPS stimulation. Both LPS-induced deficits in trans-endothelial electrical resistance (**Suppl. Fig. 1a**) and paracellular tracer permeability (**Suppl. Fig. 1b**) were fully attenuated by higher doses of propionate, without any obvious further effects beyond those seen with 1 μM of the SCFA.

Although hCMEC/D3 cells are a widely used *in vitro* model of the BBB, they are not without limitations, particularly in terms of their higher inherent permeability when compared with other non-human model systems [49]. To ensure the validity of our findings using hCMEC/D3 cells, we repeated these experiments using primary human brain microvascular endothelial cells (HBMECs). As with hCMEC/D3 cells, exposure of HBMEC monolayers for 12 h to propionate (1 μM) significantly attenuated the permeabilising effects of LPS exposure (subsequent 12 h stimulation, 50 ng/ml), in terms of both paracellular permeability to a 70 kDa FITC-conjugated dextran tracer (**Suppl Fig. 2a**) and trans-endothelial electrical resistance (**Suppl Fig. 2b**). Given this confirmation, subsequent experiments focused solely on the hCMEC/D3 cells as an *in vitro* BBB model.

Paracellular permeability and trans-endothelial electrical resistance are in large part dependent upon the integrity of inter-endothelial tight junctions [50], which are known to be disrupted following exposure to LPS [51]. We, therefore, examined the intracellular distribution of the key tight junction components occludin, claudin-5 and zona occludens-1 (ZO-1) following treatment with propionate and/or LPS. Exposure of hCMEC/D3 monolayers to propionate alone (1 μM, 24 h) had no noticeable effect on the intracellular distribution of any of the studied tight junction components, whereas treatment with LPS (50 ng/ml, 12 h) caused a marked disruption in the localisation of all three major tight junction molecules, characterised by a loss of peri-membrane immunoreactivity (**Fig. 2c**). Notably, these effects of LPS were substantially protected against by prior treatment for 12 h with 1 μM propionate.

LPS initiates a pro-inflammatory response through binding to Toll-like receptor 4, TLR4, in a complex with the accessory proteins CD14 and LY96 (MD2) [52]; we, therefore, examined expression of TLR4 signalling components as an explanation for the protective effects of propionate upon this pathway. While propionate treatment of hCMEC/D3 cells (1 μM, 24 h) had no significant effect upon expression of mRNA for TLR4 or LY96 (data not shown), such treatment significantly down-regulated expression of *CD14* mRNA (**Fig. 2d**), an effect replicated at the level of cell surface CD14 protein expression (**Fig. 2e, f**).

### NFE2L2 (NRF2) signalling and protection from oxidative stress

Enrichr (WikiPathways) analysis indicated that exposure of hCMEC/D3 cells to propionate resulted in the regulation of a number of antioxidant systems. Of known human anti-oxidant genes [53], 58 were detected on the array. We had also identified an additional 6 genes via [54] (**Supplementary Table 2**). Searches of the genes associated with each of the individual pathways referenced in **Fig. 1f** strongly indicated these changes occurred downstream of the transcription factor nuclear factor, erythroid 2 like 2 – NFE2L2 (**Fig. 3a**). Supporting this analysis, exposure of hCMEC/D3 cells for 24 h to 1 μM propionate caused a marked translocation of NFE2L2 from the cytoplasm to the nucleus (**Fig. 3b**). Functional analysis of antioxidant pathway activity was assessed by monitoring reactive oxygen species production in hCMEC/D3 cells following exposure to the mitochondrial complex I inhibitor rotenone (2.5 μM, 2 h). Preexposure of cells to 1 μM propionate for 24 h significantly attenuated the rate of fluorescent tracer accumulation, indicative of reduced levels of intracellular reactive oxygen species (**Fig. 3c**).

**Fig. 3:**
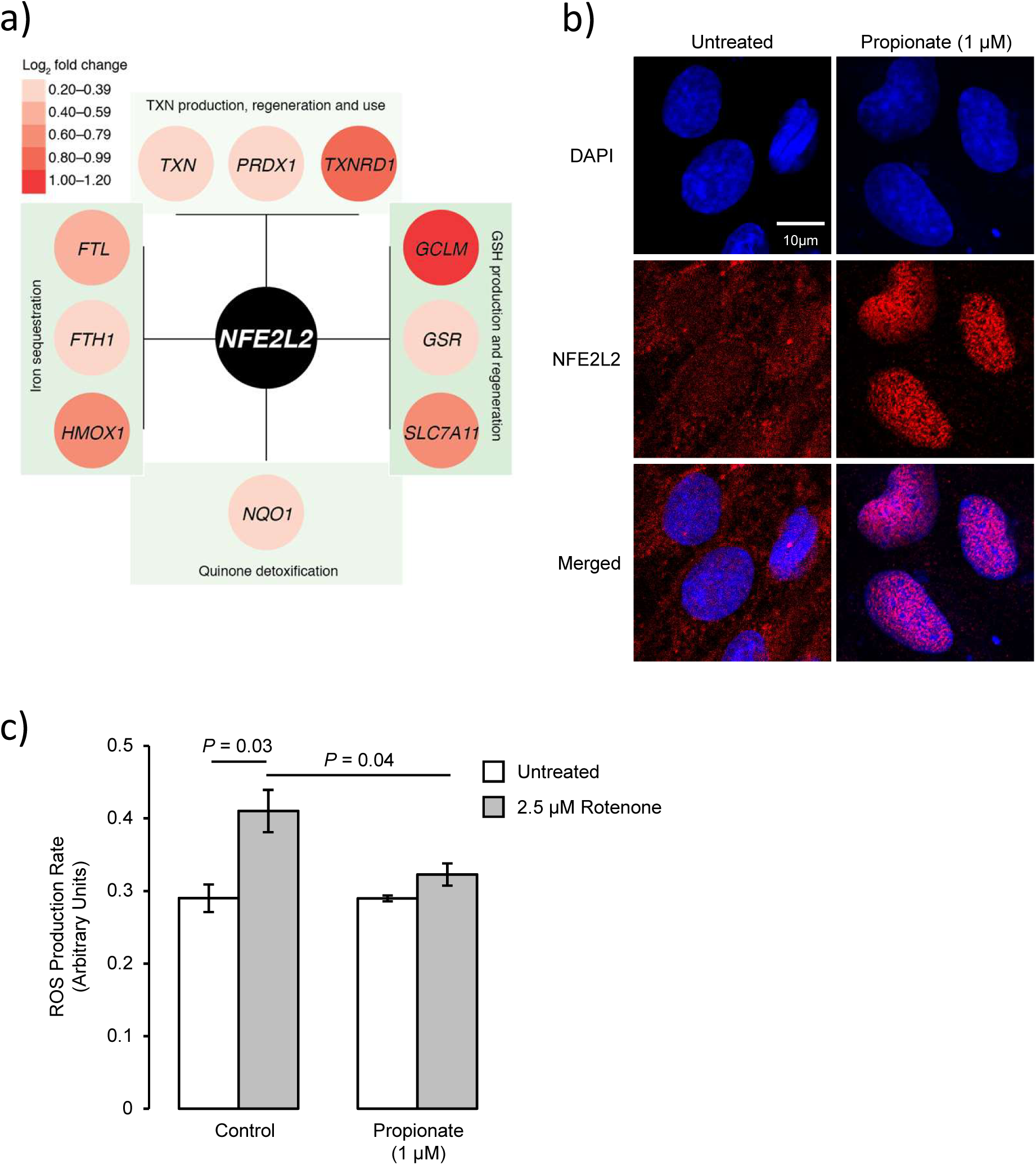
Protective effects of propionate against oxidative stress. (a) Representation of stress-response genes significantly upregulated in the current study and directly influenced by NFE2L2, *‘the master regulator of antioxidant responses’* [54]. (b) Confocal microscopic analysis of expression of NFE2L2 (Nrf2) in hCMEC/D3 cells following treatment for 24 h with 1 μM propionate; scale bar (10 μm) applies to all images. Images are representative of at least three independent experiments. (c) Production of reactive oxygen species (ROS) in control and propionate pre-treated (1 μM, 24 h) hCMEC/D3 cells treated for 30 min with the mitochondrial complex I inhibitor rotenone (2.5 μM). Data are mean ± SEM, *n*=3 independent experiments.

### Efflux transporter expression and activity

A key feature of the BBB is the expression of a wide array of efflux transporter proteins, which limit entry of numerous endogenous and xenobiotic agents to, and promote their export from, the brain. Amongst these, the proteins P-glycoprotein, BCRP and LRP-1 are prominent examples. We investigated the ability of propionate to both modify expression of these transporters and, in the case of the ABC transporter proteins *P*-glycoprotein and BCRP, serve as a direct inhibitor or substrate for the protein. Exposure of hCMEC/D3 monolayers to propionate at physiological levels (1 μM) for 24 h significantly suppressed expression of LRP-1 without modulating expression of either BCRP or P-glycoprotein (**Supplementary Fig. 1a, b**). Similarly, propionate had neither a stimulatory nor inhibitory effect upon either BCRP or P-glycoprotein activity, at concentrations between 12 nM and 27 μM (**Supplementary Fig. 1c-f**).

## Discussion

Considerable effort has gone into interrogating the gut–brain axis over recent years, with a steadily growing appreciation of the influence of the gut microbiota upon CNS function in health and disease. Mechanistic studies have identified three principal aspects to the gut-brain axis: modification of autonomic sensorimotor connections [29], immune activation [30], and regulation of neuroendocrine pathways [31], all of which incorporate a role for soluble gut-derived microbial agents, whether metabolic products or structural microbial components (e.g. LPS) themselves. In the current study, we identify a fourth facet to the gut-brain axis, namely the interactions between gut–derived microbial metabolites and the primary defensive structure of the brain, the BBB. In particular, we identify a beneficial, protective effect of the SCFA propionate upon the BBB, mitigating against deleterious inflammatory and oxidative stimuli.

If confirmed *in vivo*, our findings of protective effects of propionate upon BBB endothelial cells *in vitro* will add to the previously described beneficial actions of the SCFA upon a number of metabolic parameters. Propionate has been shown to improve glucose tolerance and insulin sensitivity, reduce high-density lipoprotein and increase serum triglyceride concentrations [35,55,56], all of which result in a more stable metabolic homeostasis. The effects of propionate upon the BBB that we describe in this study add to these pro-homeostatic actions, emphasising the contribution the SCFA plays to maintaining normal physiological function. Given that the main source of circulating propionate in humans is the intestinal microbiota [57,58], following fermentation of non-digestible carbohydrates by select bacterial species (**Fig. 4**), propionate thus represents a paradigm of commensal, mutually beneficial interactions between the host and microbiota. Moreover, consumption of food containing non-digestible carbohydrates increases circulating propionate concentrations approximately ten-fold [59,60], suggesting that the anti-inflammatory effects of the SCFA upon the cerebrovascular endothelium may be another facet of the known health benefits of high-fibre diets [61].

**Fig. 4:**
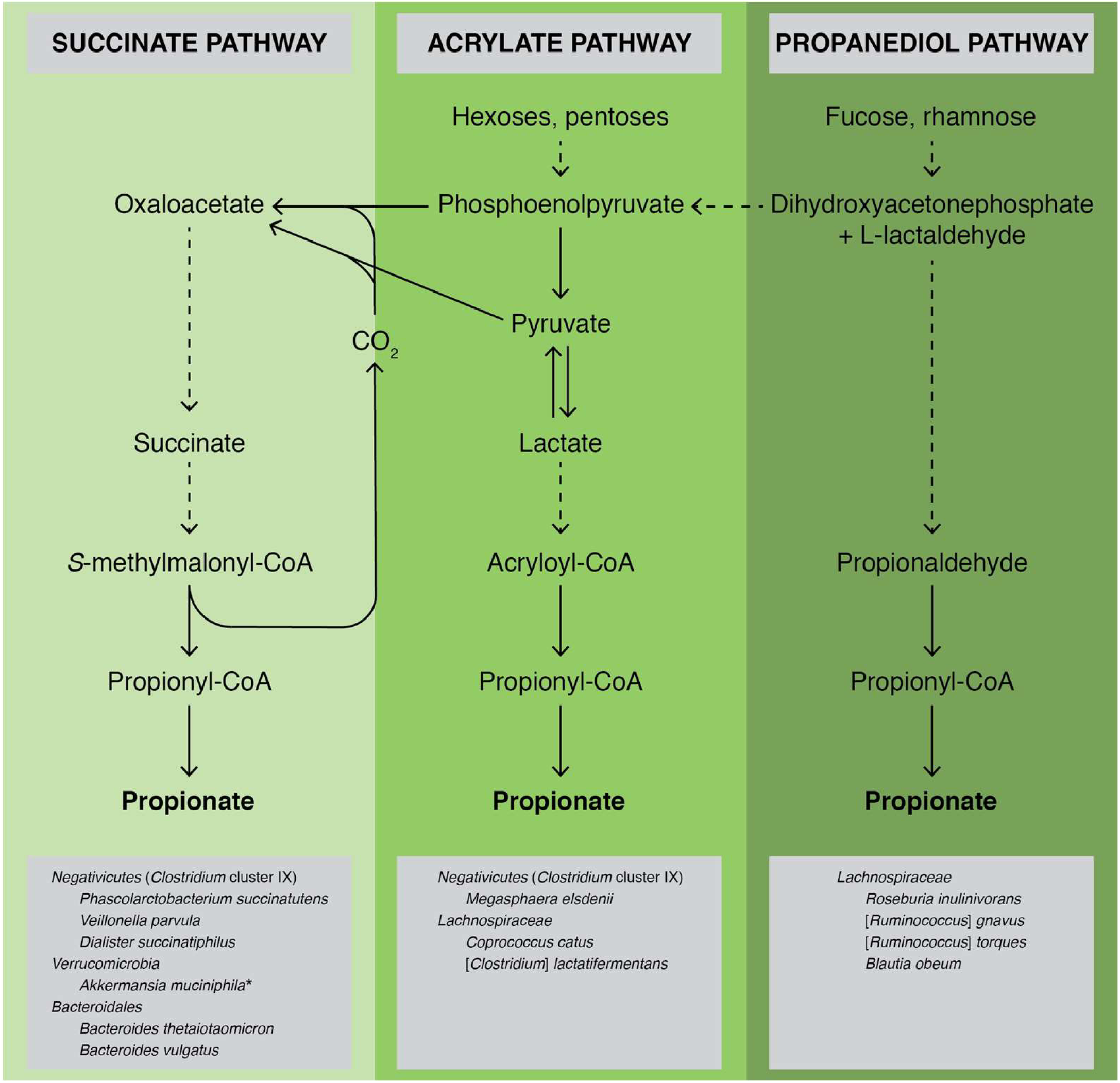
Production of propionate by the human gut microbiota. Propionate can be produced directly or indirectly by cross-feeding from succinate- and lactate-producers (e.g. *Selenomonas, Megasphaera* and *Veillonella* spp.). Image produced using information taken from [57]. \**Akkermansia muciniphila* is known to produce propionate; it is thought to do this via the succinate pathway [57].

That BBB integrity is influenced by the gut microbiota and that SCFAs may play a role in this process was recently emphasised in studies of germ-free *vs*. specific pathogen-free mice, with germ-free animals exhibiting enhanced BBB permeability and disrupted cerebral endothelial tight junctions [32]. These permeability defects were reversed fully upon conventionalisation with a pathogen-free microbiota, and partially with monocultures producing various SCFAs. Moreover, defective BBB integrity could be ameliorated at least partially by extended oral administration of sodium butyrate. Our findings thus cement SCFAs as a key group of gut-derived microbial mediators modulating BBB function, and provide evidence emphasising a direct action through the circulation. Propionate acts primarily through either of the two free fatty acid receptors FFAR2 or FFAR3 [41], which although absent from neurones in the CNS [62] have been identified in the cerebral endothelium [45], with FFAR3 confirmed herein, indicating a possible mechanism of action. Although further study would be required to prove it conclusively, our data suggest that FFAR3 may be the predominant receptor type mediating the protective effects of SCFAs, as while the major ligands for this receptor, propionate and butyrate, were both able to prevent a functional decline in BBB integrity induced by LPS exposure, this was not the case for acetate, an SCFA with greater potency at FFAR2 [39]. Future work investigating the relative contributions of the two receptor types to BBB integrity will be informative.

Notably, and perhaps unsurprisingly, SCFAs cannot fully recapitulate the BBB-restoring effects of conventionalisation of germ-free animals, as revealed in the current work and previously [32,33]. It, therefore, seems likely that additional circulating gut–derived microbial mediators may contribute to the regulation of BBB function, and are thus highly deserving of future investigation. Given that upwards of 200 distinct microbial metabolites have been identified in the circulation of healthy individuals and animals [61,63], there is clearly great potential for intestinal dysbiosis and the resultant variation in metabolite levels to influence the BBB.

This may be highly relevant to the development of neurological disease, as variation in BBB function is increasingly recognised to impact on cognitive processes, although the mechanism(s) underlying this link are poorly understood. In particular, defects in BBB integrity have been linked with impaired memory [64] and linguistic [65] function, as well as with inferior performance on psychometric tests such as the mini mental state exam [66] and Oxford handicap scale [67]. Antibiotic-induced intestinal dysbiosis has been associated with similar cognitive deficits and with a reduction in circulating gut-derived microbial metabolites [33], but as yet whether the BBB plays a role in this connection has not been investigated. If this is the case, however, as the current study suggests, regulation of BBB function by microbe-derived mediators may be an important component in some of the emerging links between intestinal dysbiosis and pathologies as significant as depression [68], Parkinson’s disease [69,70] and Alzheimer’s disease [71]. Notably, patients with early Parkinson’s or Alzheimer’s diseases have been shown to bear reduced levels of *Bacteroides* species within their faeces [71,72]. Given that *Bacteroides* spp. are important producers of SCFAs, including propionate [57], from complex carbohydrates (**Fig. 4**), this reduction may lead to a decline in circulating propionate and consequent vulnerability of the BBB, and, by extension, the brain in these major neurological conditions.

Modulatory effects of circulating gut-derived microbial metabolites upon the BBB may also be a component of the beneficial outcomes seen upon consumption of prebiotics or probiotics in a number of neurological conditions. For example, small-scale clinical trials have identified beneficial effects of probiotic drinks on cognitive ability in both Alzheimer’s disease [73] and multiple sclerosis [74], conditions associated with reduced BBB integrity [75]. Similarly, oral administration of prebiotic oligosaccharides to mice significantly reduced anxiety and stress behaviours, effects that correlated with increases in caecal acetate, propionate and butyrate concentrations [43]. Whether such changes in caecal SCFA reflected plasma levels was not measured, but given that SCFAs can be transported across the gut epithelium [76,77] increases in circulating concentrations may be likely. That inflammation contributes to depression has become clearer over recent years [78], hence it is conceivable that the anti-inflammatory effects of propionate we describe may underlie at least part of the protective effects of prebiotic treatment, a proposal which, though speculative, is deserving of further study.

In summary, we reveal here a significant new aspect of the gut-brain axis, namely the modulatory effects of circulating gut-derived microbial metabolites upon the endothelium of the BBB. Given the critical gate-keeping role the BBB plays in communication between the periphery and the brain parenchyma, our findings set the stage for future investigation of the influence the gut microbiota has on this structure, and the impact intestinal dysbiosis may have upon individual susceptibility to neurological and psychological diseases.

## Materials & Methods

### Human Tissue

Human post mortem samples were taken from the prefrontal cortex from non-neurologic controls; brains were retrieved from the UK Multiple Sclerosis Society tissue bank at Imperial College London, under ethical approval from the UK MRC Brain Bank Network (Ref. No. 08/MRE09/31+5). Brains were selected according to the following criteria: (i) availability of full clinical history, (ii) no evidence of cancer post mortem, and (iii) negligible atherosclerosis of cerebral vasculature. Tissue was fixed in 10% v/v buffered formalin and embedded in paraffin. From each paraffin block, 5 μm sections were cut and used for immunohistochemistry for FFAR3 using standard protocols [79], with a primary rabbit anti-FFAR3 polyclonal antibody (1:100; Stratech Scientific, Newmarket, UK), a horseradish peroxidase-conjugated goat anti-rabbit secondary antibody (1:300; Stratech Scientific, UK), and 2,3-diaminobenzidine and hydrogen peroxide as chromogens. Images were taken using a Leica DM5000 bright–field microscope equipped with a x40 oil immersion objective, and analysed using NIH ImageJ 1.51h (National Institutes of Health, USA).

### Cerebromicrovascular cells

The human cerebromicrovascular endothelial cell line hCMEC/D3 was purchased from VHBio Ltd (Gateshead, UK), maintained and treated as described previously [79–81]. Cells were cultured to confluency in complete EGM-2 endothelial cell growth medium (Lonza, Basel, Switzerland), whereupon medium was replaced by EGM-2 without VEGF and cells were further cultured for a minimum of 4 days to enable intercellular tight junction formation prior to experimentation. Primary human cerebromicrovascular endothelial cells (HBMEC) were purchased from Sciencell Research Laboratories (San Diego, CA, USA) and were maintained in ECM growth medium according to the supplier’s recommendations. Cells were cultured to confluency in complete ECM (Sciencell Research Laboratories, USA), whereupon medium was replaced by EGM-2 without VEGF and cells were further cultured for a minimum of 4 days to enable intercellular tight junction formation prior to experimentation. For primary cultures, trans-endothelial electrical resistance was measured as described below, and experiments were only undertaken when this had reached approximately 200 Ω.cm^2^.

### Microarrays

hCMEC/D3 cells were grown on 6-well plates coated with calf-skin collagen (Sigma-Aldrich, Gillingham, UK) to confluency as described above, further cultured for 4 days in EGM-2 medium without VEGF and exposed to propionate (1 μM, 24 h). Cells were collected into TRIzol (Thermo-Fisher Scientific, UK) and total RNA was extracted using a TRIzol Plus RNA purification kit (Thermo-Fisher Scientific, UK) and quantified using an ND-1000 Spectrophotometer (NanoDrop, Wilmington, USA).

Hybridization experiments were performed by Macrogen Inc. (Seoul, Korea) using Illumina HumanHT-12 v4.0 Expression BeadChips (Illumina Inc., San Diego, CA). RNA purity and integrity were evaluated using an ND-1000 Spectrophotometer (NanoDrop, USA) and an Agilent 2100 Bioanalyzer (Agilent Technologies, Palo Alto, USA). Total RNA was amplified and purified using TargetAmp-Nano Labelling Kit for Illumina Expression BeadChip (EPICENTRE, Madison, USA) to yield biotinylated cRNA according to the manufacturer’s instructions. Briefly, 350 ng of total RNA was reverse-transcribed to cDNA using a T7 oligo(dT) primer. Second-strand cDNA was synthesized, *in vitro*-transcribed, and labelled with biotin-NTP. After purification, the cDNA was quantified using the ND-1000 Spectrophotometer (NanoDrop, USA).

Labelled (750 ng) cDNA samples were hybridized to each beadchip for 17 h at 58 °C, according to the manufacturer’s instructions. Detection of array signal was carried out using Amersham fluorolink streptavidin-Cy3 (GE Healthcare Bio-Sciences, Little Chalfont, UK) following the bead array manual. Arrays were scanned with an Illumina bead array reader confocal scanner according to the manufacturer’s instructions. The quality of hybridization and overall chip performance were monitored by visual inspection of both internal quality control checks and the raw scanned data. Raw data were extracted using the software provided by the manufacturer (Illumina GenomeStudio v2011.1, Gene Expression Module v1.9.0).

### Processing and analyses of array data

Raw data supplied by Macrogen were quality-checked, log_2_-transformed and loess-normalized (2 iterations) using affy [82]. Probes annotated as ‘Bad’ or ‘No match’ in illuminaHumanv4.db [83] were removed from the dataset (*n* = 13,631) [84]. After this filtering step, only probes with valid Entrez identifiers (*n* = 28,979) were retained for further analyses. Entrez identifiers were matched to official gene symbols using ‘Homo_sapiens.gene_info’, downloaded from https://www.ncbi.nlm.nih.gov/guide/genes-expression/ on 14 January 2017. Average gene expression values were used for identification of differentially expressed genes. Array data have been deposited in ArrayExpress under accession number E-MTAB-5686.

Signaling Pathway Impact Analysis (SPIA) was used to identify Kyoto Encyclopedia of Genes and Genomes (KEGG) pathways activated or inhibited in hCMEC/D3 cells exposed to propionate [85]. Enrichr [47,48] was used to confirm KEGG findings (with respect to pathways, not their activation/inhibition) and to perform Gene Ontology (GO)- and WikiPathways-based analyses.

### In vitro barrier function assessments

Paracellular permeability and transendothelial electrical resistance were measured on 100 % confluent cultures polarised by growth on 24-well plate polyethylene terephthalate (PET) transwell inserts (surface area: 0.33 cm^2^, pore size: 0.4 μm; Appleton Woods, UK) coated with calf-skin collagen and fibronectin (Sigma-Aldrich, UK). The permeability of endothelial cell monolayers to 70 kDa FITC-dextran (2 mg/ml) was measured as described previously [81,86,87]; data are presented as the contribution to the permeability barrier provided by endothelial cells, Pe, throughout. Transendothelial electrical resistance (TEER) measurements were performed using a Millicell ERS-2 Voltohmmeter (Millipore, Watford, UK) and were expressed as Ω.cm^2^. In all cases, values obtained from cell-free inserts similarly coated with collagen and fibronectin were subtracted from the total values. Briefly, cells were treated with propionate (1 μM) for 24 h prior to analysis of barrier function. In some cases, barrier integrity was tested by challenge with bacterial lipopolysaccharide (LPS). Confluent endothelial monolayers were treated with propionate (1 μM) for 12 h, whereupon LPS (*Escherichia coli* O111:B4; 50 ng/ml, comparable to circulating levels of LPS in human endotoxemia [88]) was added for a further 12 h, without wash-out. Barrier function characteristics were then interrogated as described above.

### Efflux transporter assays

Activity of the major efflux transporters P-glycoprotein and BCRP [89] was determined through the use of commercially available assays (Solvo Biotechnology Inc., Budapest, Hungary), performed according to the manufacturer’s instructions. Stepwise dose-response curves centred around reported physiological circulating concentrations of propionate [90] were constructed (n=2) and both activating and inhibitory effects of propionate upon transporter activity were analysed.

### Flow cytometry analysis

hCMEC/D3 cells were labelled with APC-conjugated mouse monoclonal anti-CD14 (Thermo-Fisher Scientific, Paisley, UK), APC-conjugated mouse monoclonal anti-BCRP (BD Biosciences, Oxford, UK), FITC-conjugated mouse monoclonal LRP1 (BD Biosciences, UK), PE-conjugated mouse monoclonal anti-MDR1A (BD Biosciences, UK), unconjugated rabbit polyclonal antibody directed against FFAR3/GPR41 (Flarebio Biotech LLC, College Park, MD, USA) followed by incubation with an AF488-conjugated goat anti-rabbit secondary antibody (Thermo-Fisher Scientific, UK), or appropriate isotype controls (all BD Biosciences, UK) for analysis by flow cytometry. Briefly, hCMEC/D3 cells were treated for 24 h with propionate (1 μM), detached using 0.05 % trypsin and incubated with antibodies as described above. Immunofluorescence was analysed for 20,000 events per treatment using a BD FACSCanto II (BD Biosciences, UK) flow cytometer and data were analysed using FlowJo 8.0 software (Treestar Inc., CA, USA).

### Immunofluorescence analysis

hCMEC/D3 cells were cultured on Lab-Tek^™^ Permanox^™^ 8-well chamber slides coated with calf-skin collagen (Sigma-Aldrich, UK), prior to immunostaining according to standard protocols [79,81] and using primary antibodies directed against Nrf2 (1:500, Novus Biologicals Ltd., Abingdon, UK), occludin (1:200, Thermo-Fisher Scientific, UK), claudin-5 (1:200, Thermo-Fisher Scientific, UK) and zona occludens-1 (ZO-1; 1:100, Thermo-Fisher Scientific, UK). Nuclei were counterstained with DAPI (Sigma-Aldrich, UK). Images were captured using an LSM880 confocal laser scanning microscope (Carl Zeiss Ltd., Cambridge, UK) fitted with 405 nm, 488 nm, and 561 nm lasers, and a 63x oil immersion objective lens (NA, 1.4 mm, working distance, 0.17 mm). Images were captured with ZEN imaging software (Carl Zeiss Ltd., UK) and analysed using ImageJ 1.51h (National Institutes of Health, USA).

### Statistical analyses

Sample sizes were calculated to detect differences of 15 % or more with a power of 0.85 and α set at 5 %, calculations being informed by previously published data [79,81]. *In vitro* experimental data are expressed as mean ± SEM, with *n*=3 independent experiments performed in triplicate for all studies. In all cases, normality of distribution was established using the Shapiro–Wilkes test, followed by analysis with two-tailed Student’s t-tests to compare two groups or, for multiple comparison analysis, 1 - or 2-way ANOVA followed by Tukey’s HSD *post hoc* test. Where data was not normally distributed, non-parametric analysis was performed using the Wilcoxon signed rank test. A *P* value of less than or equal to 5 % was considered significant. Differentially expressed genes were identified in microarray data using LIMMA [91]; *P* values were corrected for multiple testing using the Benjamini–Hochberg procedure (False Discovery Rate); a *P* value of less than or equal to 10 % was considered significant in this case.

## Declarations

### Ethics approval and consent to participate

Not applicable

### Consent for publication

Not applicable

### Availability of data and material

Array data have been deposited in ArrayExpress under accession number E-MTAB-5686 (http://www.ebi.ac.uk/arrayexpress/experiments/E-MTAB-5686/)

### Authors’ contributions

LH and SM conceived the experiments; LH, TS, UU and SM performed experiments; LH and SM analysed the data; LH and SM wrote the paper; JKN, SRC and RCG provided valuable insight and advice throughout the project.

## Acknowledgements

Not applicable

## Figure Legends

**Supplementary Fig. 1:**
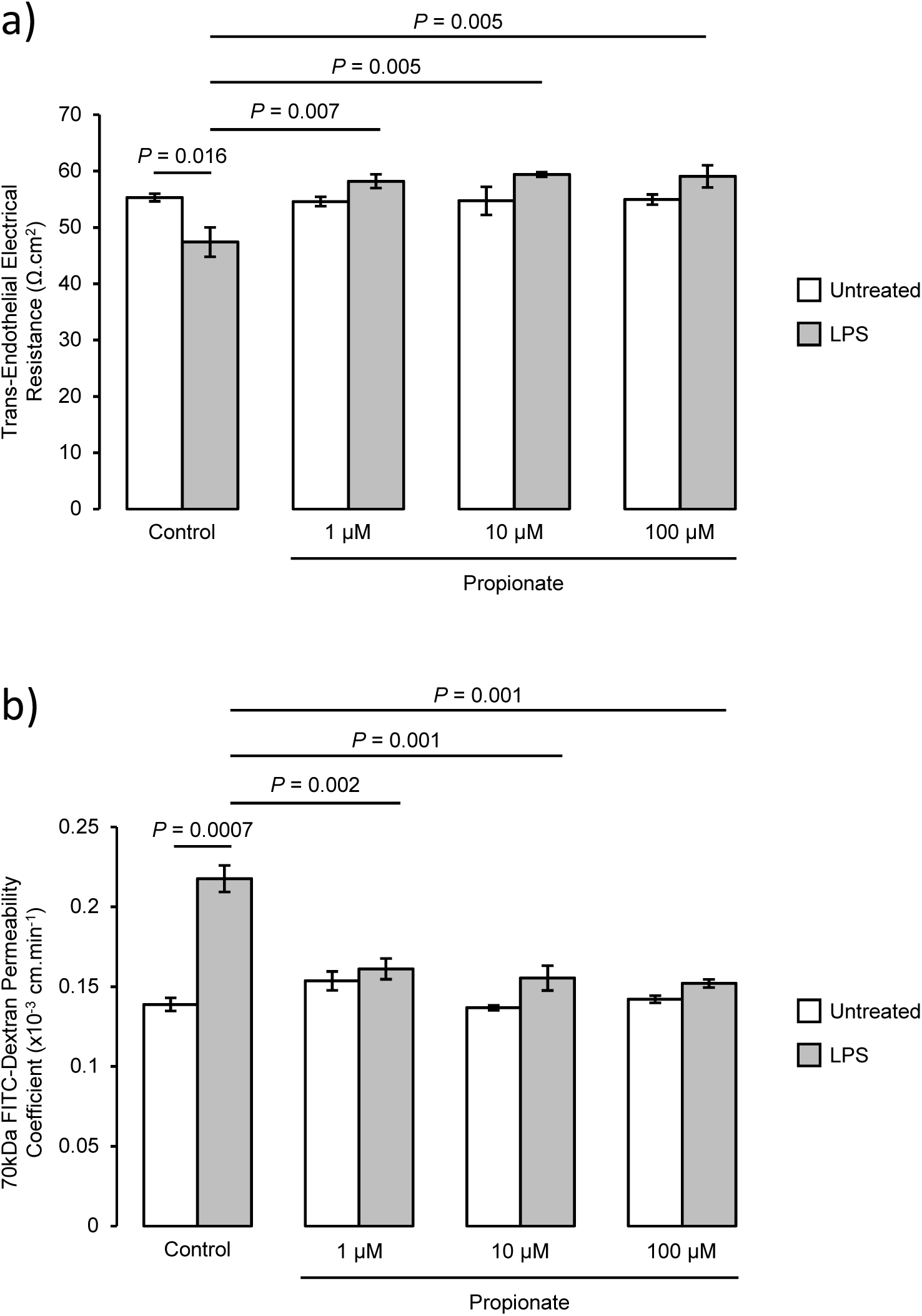
Persistence of the protective effect of propionate upon LPS-induced barrier disruption across different doses. (a) Assessment of the paracellular permeability of hCMEC/D3 monolayers to 70 kDa FITC-dextran following treatment for 24 h with 1, 10 or 100 μM propionate, with or without inclusion of 50 ng/ml LPS for the last 12 h of incubation; data are mean ± SEM, *n* = 3 independent experiments. (b) Trans-endothelial electrical resistance of hCMEC/D3 monolayers following treatment for 24 h with 1, 10 or 100 μM propionate, with or without inclusion of 50 ng/ml LPS for the last 12 h of incubation; data are mean ± SEM, *n* = 3 independent experiments.

**Supplementary Fig. 2:**
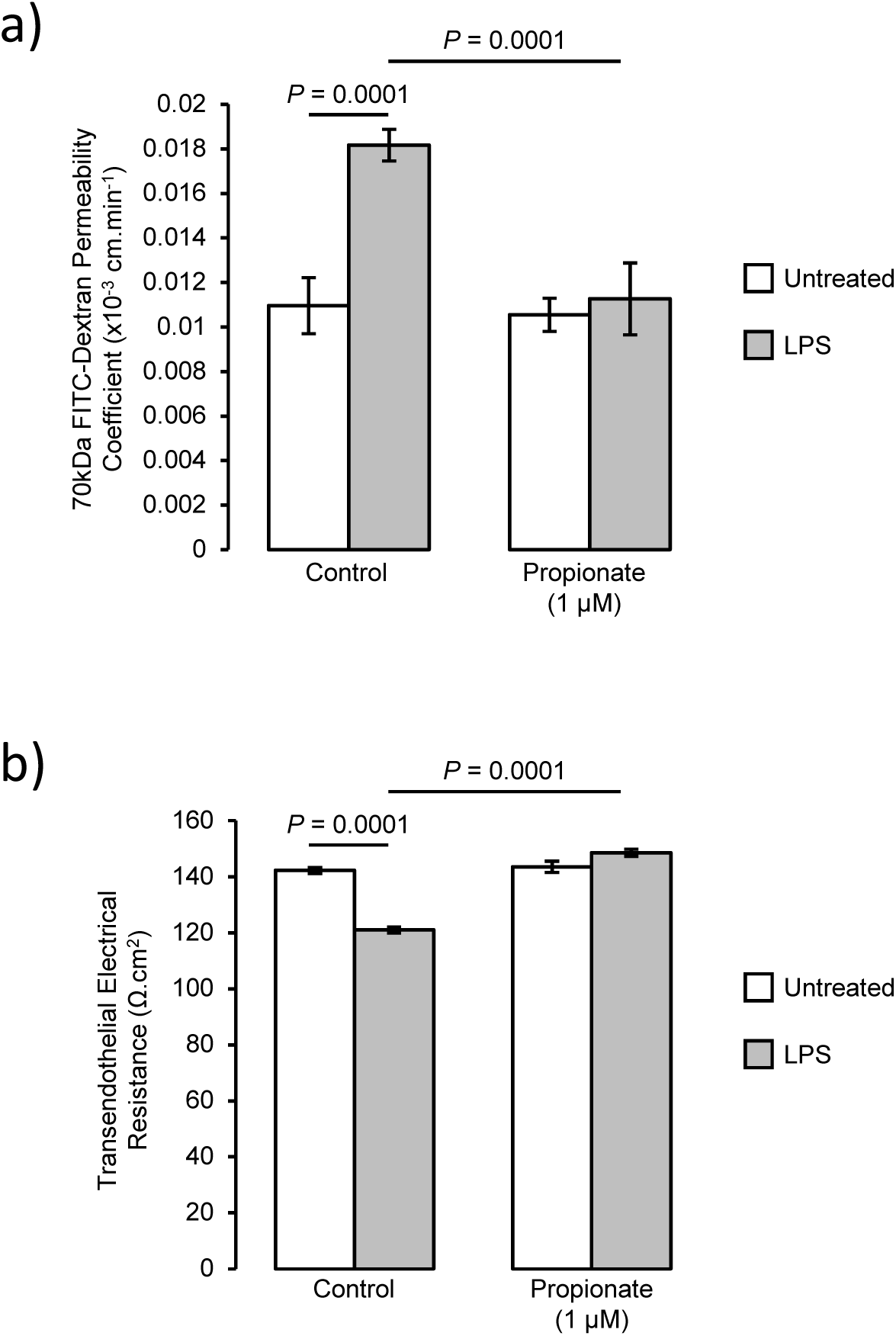
Protective effects of propionate against LPS-induced barrier disruption in primary human brain microvascular endothelial cells (HBMEC). (a) Assessment of the paracellular permeability of HBMEC monolayers to 70 kDa FITC-dextran following treatment for 24 h with 1 μM propionate, with or without inclusion of 50 ng/ml LPS for the last 12 h of incubation; data are mean ± SEM, *n* = 3 independent experiments. (b) Trans-endothelial electrical resistance of HBMEC monolayers following treatment for 24 h with 1 μM propionate, with or without inclusion of 50 ng/ml LPS for the last 12 h of incubation; data are mean ± SEM, *n* = 3 independent experiments.

**Supplementary Fig. 3:**
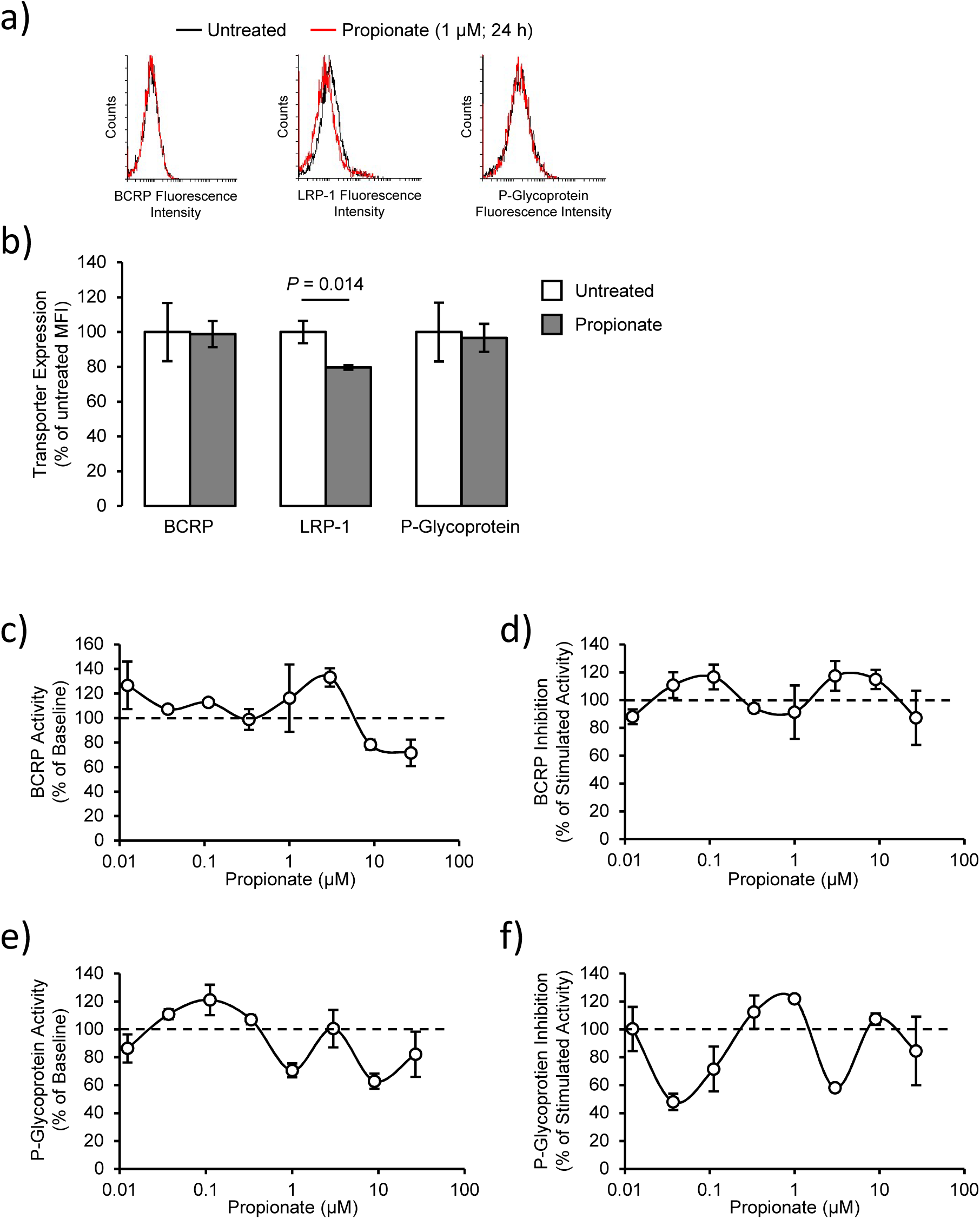
Effects of propionate upon expression and activity of typical cerebromicrovascular efflux transporter systems. (a) Surface expression of BCRP, LRP-1 and P-glycoprotein on control and propionate-treated (1 μM, 24 h) hCMEC/D3 cells (black, control, red, propionate), data are representative of three independent experiments. (b) Median fluorescence intensity of surface expression of BCRP, LRP-1 and P-glycoprotein on control and propionate-treated (1 μM, 24 h) hCMEC/D3 cells; data are mean ± SEM, n=3 independent experiments. (c) Lack of stimulatory effect of propionate upon BCRP, data are mean ± SEM, *n* = 4. (d) Lack of inhibitory effect of propionate upon stimulated ATP-dependent activity of BCRP, data are mean ± SEM, *n* = 4. (e) Lack of stimulatory effect of propionate upon P-glycoprotein, data are mean ± SEM, *n* = 4. (f) Lack of inhibitory effect of propionate upon stimulated ATP-dependent activity of P-glycoprotein, data are mean ± SEM, *n* = 4.

**Supplementary Table 1:** Effects of propionate treatment (1 μM, 24 h) upon mRNA expression of BBB-related genes in hCMEC/D3 cells, grouped in broad functional categories. Gene names listed in bold were significantly regulated compared to untreated cells (*P*_FDR_ < 0.05)

Cell Adhesion/Junctional proteins/Cytoskeletal factors
| Symbol | Description | logFC | adj.P.Val |
| --- | --- | --- | --- |
| **PECAM1** | **platelet and endothelial cell adhesion molecule 1** | **-0.518** | **0.002** |
| **CLDN11** | **claudin 11** | **0.541** | **0.024** |
| GJA1 | gap junction protein alpha 1 | 0.434 | 0.055 |
| CLDN1 | claudin 1 | 0.264 | 0.062 |
| JAM3 | junctional adhesion molecule 3 | 0.180 | 0.132 |
| UTRN | utrophin | -0.155 | 0.157 |
| CDH2 | cadherin 2 | 0.167 | 0.184 |
| CLDN7 | claudin 7 | -0.122 | 0.270 |
| ANXA1 | annexin A1 | 0.145 | 0.270 |
| TJP2 | tight junction protein 2 | -0.124 | 0.281 |
| CLDN17 | claudin 17 | -0.124 | 0.282 |
| CLDN4 | claudin 4 | 0.141 | 0.341 |
| SMARCA2 | SWI/SNF related, matrix associated, actin dependent regulator of chromatin, subfamily a, member 2 | 0.118 | 0.373 |
| CLDN23 | claudin 23 | -0.108 | 0.462 |
| JAM2 | junctional adhesion molecule 2 | -0.110 | 0.540 |
| TJP1 | tight junction protein 1 | 0.090 | 0.573 |
| CLDN6 | claudin 6 | 0.092 | 0.582 |
| LAMA4 | laminin subunit alpha 4 | -0.078 | 0.629 |
| LAMA3 | laminin subunit alpha 3 | -0.070 | 0.657 |
| DAG1 | dystroglycan 1 | -0.067 | 0.663 |
| CLDN20 | claudin 20 | 0.073 | 0.666 |
| AGRN | agrin | 0.055 | 0.694 |
| CLDN12 | claudin 12 | 0.064 | 0.751 |
| CLDN8 | claudin 8 | -0.049 | 0.753 |
| CLDN15 | claudin 15 | -0.050 | 0.778 |
| CTNNB1 | catenin beta 1 | -0.047 | 0.783 |
| VIM | vimentin | 0.041 | 0.792 |
| HAPLN2 | hyaluronan and proteoglycan link protein 2 | -0.054 | 0.794 |
| DTNA | dystrobrevin alpha | 0.053 | 0.796 |
| ESAM | endothelial cell adhesion molecule | -0.043 | 0.799 |
| LAMB2 | laminin subunit beta 2 | -0.044 | 0.803 |
| CLDN9 | claudin 9 | -0.039 | 0.804 |
| LAMA2 | laminin subunit alpha 2 | -0.057 | 0.808 |
| ITM2A | integral membrane protein 2A | -0.041 | 0.837 |
| FN1 | fibronectin 1 | -0.037 | 0.852 |
| COL4A1 | collagen type IV alpha 1 chain | 0.030 | 0.875 |
| TJP3 | tight junction protein 3 | -0.027 | 0.894 |
| CLDN3 | claudin 3 | -0.023 | 0.906 |
| GJB6 | gap junction protein beta 6 | 0.022 | 0.911 |
| CDH5 | cadherin 5 | -0.029 | 0.920 |
| LAMA1 | laminin subunit alpha 1 | 0.014 | 0.948 |
| CLDN5 | claudin 5 | -0.016 | 0.962 |
| CLDN22 | claudin 22 | 0.013 | 0.963 |
| ACTB | actin beta | 0.009 | 0.965 |
| CLDN10 | claudin 10 | -0.009 | 0.966 |
| ADGRA2 | adhesion G protein-coupled receptor A2 | 0.010 | 0.966 |
| ITGA3 | integrin subunit alpha 3 | -0.009 | 0.967 |
| OCLN | occludin | -0.007 | 0.969 |
| HSPG2 | heparan sulfate proteoglycan 2 | 0.008 | 0.973 |
| DMD | dystrophin | -0.003 | 0.989 |
| AFDN | afadin, adherens junction formation factor | 0.003 | 0.989 |
| MARVELD2 | MARVEL domain containing 2 | -0.001 | 0.996 |

Transporter proteins
| <u>Symbol</u> | <u>Description</u> | <u>logFC</u> | <u>adj.P.Val</u> |
| --- | --- | --- | --- |
| **SLC1A5** | **solute carrier family 1 member 5** | **0.400** | **0.011** |
| **SLC44A1** | **solute carrier family 44 member 1** | **-0.261** | **0.030** |
| SLC7A5 | solute carrier family 7 member 5 | 0.206 | 0.092 |
| TFRC | transferrin receptor | 0.262 | 0.099 |
| SLC38A5 | solute carrier family 38 member 5 | 0.194 | 0.165 |
| SLC38A3 | solute carrier family 38 member 3 | 0.140 | 0.240 |
| SLC22A5 | solute carrier family 22 member 5 | 0.140 | 0.272 |
| SLC29A4 | solute carrier family 29 member 4 | 0.144 | 0.299 |
| SLC22A8 | solute carrier family 22 member 8 | -0.126 | 0.308 |
| SLC2A1 | solute carrier family 2 member 1 | -0.129 | 0.342 |
| SLC38A2 | solute carrier family 38 member 2 | 0.120 | 0.381 |
| SLC28A2 | solute carrier family 28 member 2 | 0.133 | 0.411 |
| SLC5A1 | solute carrier family 5 member 1 | 0.100 | 0.447 |
| SLC5A6 | solute carrier family 5 member 6 | 0.094 | 0.452 |
| SLC6A6 | solute carrier family 6 member 6 | 0.096 | 0.461 |
| SLC1A4 | solute carrier family 1 member 4 | -0.115 | 0.462 |
| SLC27A4 | solute carrier family 27 member 4 | -0.139 | 0.463 |
| LRP2 | LDL receptor related protein 2 | 0.082 | 0.501 |
| SLC38A1 | solute carrier family 38 member 1 | 0.091 | 0.510 |
| SLC22A1 | solute carrier family 22 member 1 | 0.078 | 0.560 |
| LDLR | low density lipoprotein receptor | -0.075 | 0.566 |
| SLC1A3 | solute carrier family 1 member 3 | 0.084 | 0.581 |
| MFSD2A | major facilitator superfamily domain containing 2A | 0.079 | 0.593 |
| ABCG2 | ATP binding cassette subfamily G member 2 (Junior blood group) | 0.066 | 0.671 |
| INSR | insulin receptor | 0.060 | 0.718 |
| AQP4 | aquaporin 4 | 0.060 | 0.733 |
| SLC16A2 | solute carrier family 16 member 2 | -0.057 | 0.780 |
| ABCC5 | ATP binding cassette subfamily C member 5 | -0.041 | 0.793 |
| SLCO1C1 | solute carrier organic anion transporter family member 1C1 | 0.041 | 0.795 |
| SLC29A1 | solute carrier family 29 member 1 (Augustine blood group) | 0.036 | 0.807 |
| SLC27A1 | solute carrier family 27 member 1 | -0.036 | 0.818 |
| SLC7A3 | solute carrier family 7 member 3 | 0.038 | 0.824 |
| SLC22A2 | solute carrier family 22 member 2 | 0.035 | 0.843 |
| SLC16A1 | solute carrier family 16 member 1 | -0.047 | 0.847 |
| ABCB1 | ATP binding cassette subfamily B member 1 | 0.029 | 0.866 |
| AGER | advanced glycosylation end-product specific receptor | -0.026 | 0.908 |
| AVPR1A | arginine vasopressin receptor 1A | -0.023 | 0.912 |
| ABCA2 | ATP binding cassette subfamily A member 2 | 0.015 | 0.947 |
| SLC6A9 | solute carrier family 6 member 9 | 0.013 | 0.949 |
| SLC1A1 | solute carrier family 1 member 1 | -0.013 | 0.954 |
| SLC7A1 | solute carrier family 7 member 1 | 0.013 | 0.955 |
| ABCC1 | ATP binding cassette subfamily C member 1 | 0.012 | 0.956 |
| SLC22A3 | solute carrier family 22 member 3 | -0.012 | 0.957 |
| LEPR | leptin receptor | -0.009 | 0.960 |
| SLC16A7 | solute carrier family 16 member 7 | -0.012 | 0.962 |
| ABCC4 | ATP binding cassette subfamily C member 4 | -0.011 | 0.963 |
| SLC5A3 | solute carrier family 5 member 3 | -0.009 | 0.967 |
| SLC7A6 | solute carrier family 7 member 6 | -0.008 | 0.969 |
| SLCO2B1 | solute carrier organic anion transporter family member 2B1 | -0.005 | 0.983 |
| ABCC2 | ATP binding cassette subfamily C member 2 | -0.003 | 0.988 |
| SLCO1B1 | solute carrier organic anion transporter family member 1B1 | 0.003 | 0.988 |
| SLC2A13 | solute carrier family 2 member 13 | 0.003 | 0.990 |
| SLC1A2 | solute carrier family 1 member 2 | 0.001 | 0.995 |

Inflammatory response
| <u>Symbol</u> | <u>Description</u> | <u>logFC</u> | <u>adj.P.Val</u> |
| --- | --- | --- | --- |
| **TNFSF10** | **tumor necrosis factor superfamily member 10** | **-0.684** | **0.001** |
| **PDGFRB** | **platelet derived growth factor receptor beta** | **-0.441** | **0.015** |
| **TNFRSF1A** | **TNF receptor superfamily member 1A** | **-0.289** | **0.021** |
| **TNFRSF12A** | **TNF receptor superfamily member 12A** | **0.383** | **0.028** |
| **TNFRSF21** | **TNF receptor superfamily member 21** | **0.325** | **0.031** |
| ITGB4 | integrin subunit beta 4 | -0.205 | 0.056 |
| TNFAIP6 | TNF alpha induced protein 6 | 0.325 | 0.118 |
| PODXL | podocalyxin like | -0.194 | 0.130 |
| ITGA5 | integrin subunit alpha 5 | -0.211 | 0.163 |
| ITGA1 | integrin subunit alpha 1 | -0.150 | 0.188 |
| PTGS2 | prostaglandin-endoperoxide synthase 2 | 0.187 | 0.189 |
| ITGB5 | integrin subunit beta 5 | -0.156 | 0.193 |
| CXCL2 | C-X-C motif chemokine ligand 2 | 0.171 | 0.231 |
| IKBKB | inhibitor of nuclear factor kappa B kinase subunit beta | -0.139 | 0.299 |
| SOD1 | superoxide dismutase 1, soluble | 0.126 | 0.338 |
| ITGB8 | integrin subunit beta 8 | -0.144 | 0.340 |
| NOS1 | nitric oxide synthase 1 | 0.114 | 0.366 |
| CCR5 | C-C motif chemokine receptor 5 (gene/pseudogene) | 0.222 | 0.391 |
| ITGA4 | integrin subunit alpha 4 | 0.161 | 0.430 |
| CLEC5A | C-type lectin domain family 5 member A | 0.138 | 0.441 |
| ITGA6 | integrin subunit alpha 6 | -0.092 | 0.442 |
| GRN | granulin precursor | -0.089 | 0.455 |
| MMP9 | matrix metalloproteinase 9 | -0.099 | 0.475 |
| NR3C1 | nuclear receptor subfamily 3 group C member 1 | -0.085 | 0.496 |
| CRH | corticotropin releasing hormone | -0.092 | 0.558 |
| AGT | angiotensinogen | -0.091 | 0.594 |
| PTGDS | prostaglandin D2 synthase | -0.097 | 0.596 |
| NOX4 | NADPH oxidase 4 | 0.070 | 0.601 |
| MMP2 | matrix metalloproteinase 2 | -0.088 | 0.687 |
| SELP | selectin P | -0.074 | 0.689 |
| IL1RN | interleukin 1 receptor antagonist | 0.060 | 0.692 |
| CXCR3 | C-X-C motif chemokine receptor 3 | -0.060 | 0.711 |
| F11R | F11 receptor | -0.089 | 0.741 |
| TNFRSF1B | TNF receptor superfamily member 1B | -0.098 | 0.760 |
| SEMA7A | semaphorin 7A (John Milton Hagen blood group) | 0.054 | 0.770 |
| ITGB3 | integrin subunit beta 3 | 0.049 | 0.793 |
| ITGAV | integrin subunit alpha V | -0.033 | 0.837 |
| TLR2 | toll like receptor 2 | 0.030 | 0.860 |
| ITGB1 | integrin subunit beta 1 | 0.026 | 0.880 |
| PTGER3 | prostaglandin E receptor 3 | -0.022 | 0.895 |
| TNF | tumor necrosis factor | -0.016 | 0.927 |
| ITGB2 | integrin subunit beta 2 | -0.012 | 0.951 |
| IL1B | interleukin 1 beta | -0.033 | 0.967 |
| CCR2 | C-C motif chemokine receptor 2 | -0.007 | 0.970 |
| CD276 | CD276 molecule | 0.006 | 0.973 |
| C3 | complement C3 | -0.001 | 0.997 |

Vascular function/coagulation cascade
| <u>Symbol</u> | <u>Description</u> | <u>logFC</u> | <u>adj.P.Val</u> |
| --- | --- | --- | --- |
| **SERPINE2** | **serpin family E member 2** | **0.461** | **0.007** |
| **PROCR** | **protein C receptor** | **0.240** | **0.046** |
| PLAT | plasminogen activator, tissue type | -0.242 | 0.051 |
| SERPINE1 | serpin family E member 1 | 0.244 | 0.212 |
| PROS1 | protein S (alpha) | -0.165 | 0.262 |
| PROC | protein C, inactivator of coagulation factors Va and VIIIa | -0.128 | 0.479 |
| CA1 | carbonic anhydrase 1 | 0.081 | 0.518 |
| VWF | von Willebrand factor | -0.139 | 0.567 |
| AVP | arginine vasopressin | -0.057 | 0.758 |
| SERPINI1 | serpin family I member 1 | -0.033 | 0.840 |
| PLG | plasminogen | -0.030 | 0.884 |
| KNG1 | kininogen 1 | -0.024 | 0.898 |
| NOS3 | nitric oxide synthase 3 | 0.037 | 0.905 |
| MYLK | myosin light chain kinase | -0.014 | 0.949 |
| PTAFR | platelet activating factor receptor | -0.013 | 0.952 |
| EPAS1 | endothelial PAS domain protein 1 | 0.010 | 0.955 |

Endothelial proliferation/angiogenesis
| <u>Symbol</u> | <u>Description</u> | <u>logFC</u> | <u>adj.P.Val</u> |
| --- | --- | --- | --- |
| PDGFB | platelet derived growth factor subunit B | -0.226 | 0.090 |
| TMEFF2 | transmembrane protein with EGF like and two follistatin like domains 2 | 0.166 | 0.170 |
| S100A12 | S100 calcium binding protein A12 | 0.143 | 0.245 |
| FGF19 | fibroblast growth factor 19 | -0.157 | 0.354 |
| IGFBP3 | insulin like growth factor binding protein 3 | 0.091 | 0.486 |
| RGS5 | regulator of G-protein signaling 5 | -0.075 | 0.548 |
| FLT1 | fms related tyrosine kinase 1 | -0.112 | 0.572 |
| HNRNPDL | heterogeneous nuclear ribonucleoprotein D like | -0.084 | 0.601 |
| VEGFA | vascular endothelial growth factor A | -0.072 | 0.617 |
| S100B | S100 calcium binding protein B | -0.071 | 0.643 |
| EZH1 | enhancer of zeste 1 polycomb repressive complex 2 subunit | -0.068 | 0.722 |
| PTPRB | protein tyrosine phosphatase, receptor type B | -0.057 | 0.745 |
| HMGB1 | high mobility group box 1 | 0.044 | 0.775 |
| PTN | pleiotrophin | -0.029 | 0.920 |
| KDR | kinase insert domain receptor | 0.022 | 0.934 |
| BTG2 | BTG anti-proliferation factor 2 | 0.012 | 0.958 |
| EPO | erythropoietin | -0.011 | 0.963 |

Other BBB-related genes
| <u>Symbol</u> | <u>Description</u> | <u>logFC</u> | <u>adj.P.Val</u> |
| --- | --- | --- | --- |
| EPHA2 | EPH receptor A2 | -0.249 | 0.076 |
| MOG | myelin oligodendrocyte glycoprotein | -0.090 | 0.546 |
| CLN3 | CLN3, battenin | -0.085 | 0.566 |
| SRGN | serglycin | 0.072 | 0.574 |
| MBP | myelin basic protein | -0.066 | 0.646 |
| RAMP2 | receptor activity modifying protein 2 | 0.054 | 0.713 |
| CLCN2 | chloride voltage-gated channel 2 | -0.055 | 0.733 |
| CPE | carboxypeptidase E | 0.044 | 0.811 |
| CYBB | cytochrome b-245 beta chain | 0.033 | 0.856 |
| MPZL1 | myelin protein zero like 1 | 0.028 | 0.864 |
| GAB2 | GRB2 associated binding protein 2 | -0.030 | 0.866 |
| MAP3K7 | mitogen-activated protein kinase kinase kinase 7 | 0.028 | 0.882 |
| APP | amyloid beta precursor protein | -0.044 | 0.890 |
| APLP2 | amyloid beta precursor like protein 2 | 0.022 | 0.914 |
| PLP1 | proteolipid protein 1 | 0.019 | 0.935 |
| HDC | histidine decarboxylase | 0.007 | 0.985 |
| HRH3 | histamine receptor H3 | 0.003 | 0.989 |
| APOE | apolipoprotein E | 0.000 | 1.000 |
| GFAP | glial fibrillary acidic protein | 0.000 | 1.000 |

**Supplementary Table 2:** Effects of propionate treatment (1 μM, 24 h) upon mRNA expression of antioxidant system-related genes in hCMEC/D3 cells. Gene names listed in bold were significantly regulated compared to untreated cells (*P*_FDR_ < 0.05).

| Symbol | log<sub>2</sub> fold change | Adjusted *P* value | Synonyms | Description |
| --- | --- | --- | --- | --- |
| GCLM† | 1.034 | 1.312×10<sup>-4</sup> | GLCLR | glutamate-cysteine ligase modifier subunit |
| SRXN1 | 1.242 | 1.934×10<sup>-4</sup> | C20orf139, Npn3, SRX, SRX1 | sulfiredoxin 1 |
| TXNRD1* | 0.928 | 2.770×10<sup>-4</sup> | GRIM-12, TR, TR1, TRXR1, TXNR | thioredoxin reductase 1 |
| HMOX1† | 0.693 | 2.770×10<sup>-4</sup> | HMOX1D, HO-1, HSP32, bK286B10 | heme oxygenase 1 |
| FTL† | 0.564 | 2.467×10<sup>-3</sup> | LFTD, NBIA3 | ferritin light chain |
| SLC7A11† | 0.649 | 3.676×10<sup>-3</sup> | CCBR1, xCT | solute carrier family 7 member 11 |
| TXNL4B | 0.388 | 6.601×10<sup>-3</sup> | DLP, Dim2 | thioredoxin like 4B |
| NQO1† | 0.345 | 0.015 | DHQU, DIA4, DTD, NMOR1, NMORI, QR1 | NAD(P)H quinone dehydrogenase 1 |
| TXNDC9 | 0.346 | 0.020 | APACD, PHLP3 | thioredoxin domain containing 9 |
| PRDX1* | 0.306 | 0.021 | MSP23, NKEF-A, NKEFA, PAG, PAGA, PAGB, PRX1, PRXI, TDPX2 | peroxiredoxin 1 |
| MT1F | -0.254 | 0.039 | MT1 | metallothionein 1F |
| MT1G | 0.224 | 0.043 | MT1, MT1K | metallothionein 1G |
| GLRX3 | 0.256 | 0.045 | GLRX4, GRX3, GRX4, PICOT, TXNL2, TXNL3 | glutaredoxin 3 |
| TXN* | 0.240 | 0.050 | TRDX, TRX, TRX1 | thioredoxin |
| FTH1† | 0.206 | 0.061 | FHC, FTH, FTHL6, HFE5, PIG15, PLIF | ferritin heavy chain 1 |
| GSR* | 0.328 | 0.061 | HEL-75, HEL-S-122m | glutathione-disulfide reductase |
| MSRA | -0.222 | 0.082 | PMSR | methionine sulfoxide reductase A |
| TXNDC5 | 0.217 | 0.085 | ENDOPDI, ERP46, HCC-2, HCC2, PDIA15, STRF8, UNQ364 | thioredoxin domain containing 5 |
| MT1M | -0.213 | 0.106 | MT-1M, MT-IM, MT1, MT1K | metallothionein 1M |
| GPX7 | 0.210 | 0.110 | CL683, GPX6, GPx-7, GSHPx-7, NPGPx | glutathione peroxidase 7 |
| TXNRD2 | -0.166 | 0.164 | SELZ, TR, TR-BETA, TR3, TRXR2 | thioredoxin reductase 2 |
| ERP44 | 0.170 | 0.165 | PDIA10, TXNDC4 | endoplasmic reticulum protein 44 |
| PRDX4 | 0.153 | 0.173 | AOE37-2, AOE372, HEL-S-97n, PRX-4 | peroxiredoxin 4 |
| SOD2 | 0.191 | 0.238 | IPO-B, IPOB, MNSOD, MVCD6, Mn-SOD | superoxide dismutase 2, mitochondrial |
| PDIA6 | 0.132 | 0.250 | ERP5, P5, TXNDC7 | protein disulfide isomerase family A member 6 |
| TXNDC8 | -0.176 | 0.262 | SPTRX-3, TRX6, bA427L11.2 | thioredoxin domain containing 8 |
| GPX4 | -0.125 | 0.315 | GPx-4, GSHPx-4, MCSP, PHGPx, SMDS, snGPx, snPHGPx | glutathione peroxidase 4 |
| SOD1 | 0.126 | 0.338 | ALS, ALS1, HEL-S-44, IPOA, SOD, hSod1, homodimer | superoxide dismutase 1, soluble |
| TMX1 | 0.140 | 0.354 | PDIA11, TMX, TXNDC, TXNDC1 | thioredoxin related transmembrane protein 1 |
| GLRX | -0.110 | 0.375 | GRX, GRX1 | glutaredoxin |
| TXNDC17 | 0.111 | 0.404 | TRP14, TXNL5 | thioredoxin domain containing 17 |
| MT1A | -0.096 | 0.479 | MT1, MT1S, MTC | metallothionein 1A |
| PRDX3 | 0.084 | 0.486 | AOP-1, AOP1, HBC189, MER5, PRO1748, SP-22, prx-III | peroxiredoxin 3 |
| GLRX2 | 0.086 | 0.500 | CGI-133, GRX2 | glutaredoxin 2 |
| NME9 | 0.093 | 0.506 | NM23-H9, TXL-2, TXL2, TXNDC6 | NME/NM23 family member 9 |
| TXNDC12 | 0.114 | 0.539 | AG1, AGR1, ERP16, ERP18, ERP19, PDIA16, TLP19, hAG-1, hTLP19 | thioredoxin domain containing 12 |
| MT1X | 0.104 | 0.544 | MT-1I, MT1 | metallothionein 1X |
| PRDX6 | 0.081 | 0.549 | 1-Cys, AOP2, HEL-S-128m, NSGPx, PRX, aiPLA2, p29 | peroxiredoxin 6 |
| GPX6 | -0.087 | 0.567 | GPX5p, GPXP3, GPx-6, GSHPx-6, dJ1186N24, dJ1186N24.1 | glutathione peroxidase 6 |
| GPX3 | -0.073 | 0.622 | GPx-P, GSHPx-3, GSHPx-P | glutathione peroxidase 3 |
| TMX2 | 0.063 | 0.637 | CGI-31, PDIA12, PIG26, TXNDC14 | thioredoxin related transmembrane protein 2 |
| CAT | 0.068 | 0.642 | - | catalase |
| PRDX5 | -0.064 | 0.662 | ACR1, AOEB166, B166, HEL-S-55, PLP, PMP20, PRDX6, PRXV, SBBI10, prx-V | peroxiredoxin 5 |
| MT1E | -0.091 | 0.687 | MT-1E, MT-IE, MT1, MTD | metallothionein 1E |
| GPX5 | 0.053 | 0.694 | HEL-S-75p | glutathione peroxidase 5 |
| GPX1 | -0.059 | 0.698 | GPXD, GSHPX1 | glutathione peroxidase 1 |
| SOD3 | -0.048 | 0.763 | EC-SOD | superoxide dismutase 3, extracellular |
| GLRX5 | -0.047 | 0.777 | C14orf87, FLB4739, GRX5, PR01238, PRO1238, PRSA, SIDBA3, SPAHGC | glutaredoxin 5 |
| SELENOP | 0.040 | 0.794 | SELP, SEPP, SEPP1, SeP | selenoprotein P |
| TXN2 | 0.052 | 0.832 | COXPD29, MT-TRX, MTRX, TRX2 | thioredoxin 2 |
| CCS | -0.031 | 0.841 | - | copper chaperone for superoxide dismutase |
| MT1H | 0.041 | 0.853 | MT-0, MT-1H, MT-IH, MT1 | metallothionein 1H |
| CP | 0.023 | 0.889 | CP-2 | ceruloplasmin |
| TXNIP | 0.151 | 0.899 | ARRDC6, EST01027, HHCPA78, THIF, VDUP1 | thioredoxin interacting protein |
| TXNDC11 | -0.021 | 0.907 | EFP1 | thioredoxin domain containing 11 |
| PRDX2 | 0.016 | 0.935 | HEL-S-2a, NKEF-B, NKEFB, PRP, PRX2, PRXII, PTX1, TDPX1, TPX1, TSA | peroxiredoxin 2 |
| MT2A | -0.014 | 0.942 | MT2 | metallothionein 2A |
| NME8 | -0.015 | 0.943 | CILD6, HEL-S-99, NM23-H8, SPTRX2, TXNDC3, sptrx-2 | NME/NM23 family member 8 |
| TXNDC2 | -0.012 | 0.953 | SPTRX, SPTRX1 | thioredoxin domain containing 2 |
| TMX4 | -0.011 | 0.961 | DJ971N18.2, PDIA14, TXNDC13 | thioredoxin related transmembrane protein 4 |
| MT1B | -0.009 | 0.963 | MT-1B, MT-IB, MT1, MT1Q, MTP | metallothionein 1B |
| TXNL1 | -0.013 | 0.968 | HEL-S-114, TRP32, TXL-1, TXNL, TxI | thioredoxin like 1 |
| GPX2 | -0.008 | 0.969 | GI-GPx, GPRP, GPRP-2, GPx-2, GPx-GI, GSHPX-GI, GSHPx-2 | glutathione peroxidase 2 |
| TMX3 | -0.007 | 0.982 | PDIA13, TXNDC10 | thioredoxin related transmembrane protein 3 |

†Anti-oxidant genes identified from Enrichr search and Gorrini *et al.* (2013), and included in **Fig. 3a**.
\*Anti-oxidant genes identified from Enrichr search, Gorrini *et al.* (2013) and Gelain *et al.* (2009), and included in **Fig. 3a**.

